# Three-dimensional fruit growth analysis clarifies developmental mechanisms underlying complex shape diversity in persimmon fruit

**DOI:** 10.1101/2023.02.07.527529

**Authors:** Akane Kusumi, Soichiro Nishiyama, Ryutaro Tao

## Abstract

How fruit size and shape are determined is of research interest in agriculture and developmental biology. Fruit typically exhibits three-dimensional structures with genotype-dependent geometric features. Although minor developmental variations have been recognized, little research has fully visualized and measured these variations throughout fruit growth. In this study, a high-resolution 3D scanner was used to investigate the fruit development of 51 persimmon (*Diospyros kaki*) cultivars with various complex shapes. We obtained 2,380 3D fruit models that fully represented fruit appearance, and enabled precise and automated measurements of unique geometric features throughout fruit development. The 3D fruit model analysis identified key stages that determined the shape attributes at maturity. Typically, genetic diversity in vertical groove development was found, and such grooves can be filled by tissue expansion in the carpal fusion zone during fruit development. Furthermore, transcriptome analysis of fruit tissues from groove/non-groove tissues revealed gene co-expression networks that were highly associated with groove depth variation. The presence of *YABBY* homologs was most closely associated with groove depth and indicated the possibility that this pathway is a key molecular contributor to vertical groove depth variation. These results demonstrate the validity of fruit 3D growth analysis, which is a powerful tool for identifying the developmental mechanisms of fruit shape variation and the molecular basis of this diversity.

## Introduction

Fruit is an organ of angiosperms that facilitates reproduction. Generally, the fruit ovary protects seeds until they mature and promotes seed dispersal once fruit has reached maturity by color or aroma to attract animals (Howe and Smallwood, 1982). Fruits are classified by three characteristics: dehiscence or indehiscence, dry or fleshy, and unfused or fused carpels (Seymour et al. 2013). Fruit with fused carpels evolved from organs that are homologous to leaves (Scutt et al., 2006). Many studies demonstrated that mutations of genes related to floral development resulted in conversion of flowers into leaves, and that ectopic expression of those genes could transform leaves into floral organs and vice versa (Ferrándiz et al., 2010; Honma and Goto, 2001). After floral organ development is completed, fertilization stimulates hormonal and genetic change in ovules and ovaries; thus, fruit development is triggered (Mezzetti et al., 2004; Eklund et al., 2010). Fleshy fruit development is typically driven by cell division at the early stage and subsequently there is cell expansion (Gillaspy et al., 1993). Several major genes have been identified as affecting the final fruit size or shape by regulating the speed and direction of cell division and expansion (van der Knaap et al., 2014).

Fruit shape is an important agricultural trait and has been targeted in breeding (Cliff et al., 2002). In agricultural research practices, fruit shape has been evaluated by descriptive rating scales or 2D imaging of fruit. In particular, 2D imaging has been used for many species with various fruit shapes, such as tomato, cucumber, apple, and litchi (Gonzalo et al., 2009; Shimomura et al., 2016; Currie et al., 2000; Osako et al., 2020). In most cases, 2D imaging methods acquire images from one side of the fruit or from a sliced cross section, and shape features are obtained by image processing. Although 2D imaging-based quantitative evaluation of fruit shape is advantageous because the data collection can be quick, inexpensive, and simple, there are also disadvantages. Ding et al. (2000) found that the biggest disadvantage of 2D fruit morphology evaluation is that there is missing or reduced information. Two-dimensional evaluation often fails to capture part of the shape features that the original 3D structure possesses in that it represents only a single side of fruit and thus important information can be obscured. For this reason, 3D plant phenotyping has emerged and been applied in some important crops such as potato and strawberry (Liu et al., 2021; He et al., 2017; Li et al., 2020). Recent advances in photogrammetry have facilitated 3D image acquisition, yet very few studies have employed this technology to assess genetic diversity in fruit development.

Cultivated oriental persimmon (*Diospyros kaki* Thunb.), a temperate deciduous fruit tree, shows unique genetic variations in fruit development and shape (Nishiyama and Yamane, 2022). Regarding overall shape, some are flat, such as ‘Hiratanenashi,’ but others are elongated, such as ‘Fudegaki.’ In addition, some cultivars develop vertical and horizontal grooves (“Mizo” and “Za” in Japanese) in the lateral side and the proximal (calyx) end of the fruit, respectively, adding unique diversity to persimmon fruit shape (Fig. 1a and 1b). Conventionally, persimmon fruit shape diversity has been categorized into phenotype classes (Fruit Tree Experiment Station of Hiroshima Prefecture, 1979; UPOV, 2004). For example, vertical groove depth was evaluated using a score and descriptor, such as “7. deep,” “3. shallow, but obvious,” or “1. absent” in the report by Fruit Tree Experiment Station of Hiroshima Prefecture (1979). Similarly, horizontal grooves were evaluated as “9. extremely big,” “3. small,” or “1. absent.” However, this qualitative categorization can produce confusion. In Fig. 1a, ‘Mizushimagosho’ and ‘Amayotsumizo’ were classified as having vertical grooves at the same level, though the actual depths are quite different. Therefore, developing a quantitative measurement method for persimmon fruit is beneficial in both agriculture and fruit development research.

**Figure 1.**
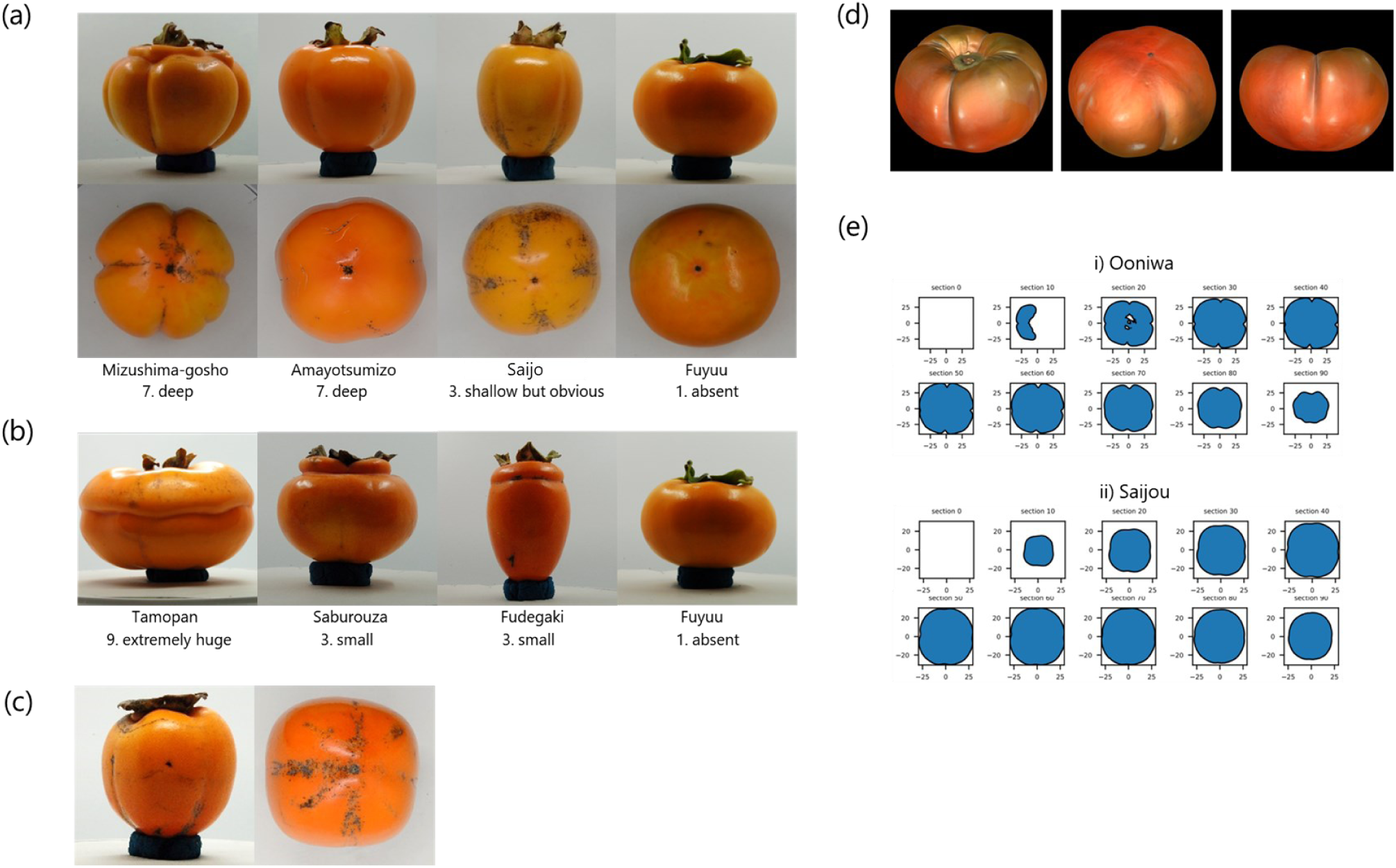
Three-dimensional scanning of fruit from diverse persimmon cultivars. (a, b) Diversity of (a) vertical and (b) horizontal grooves in persimmon fruits. (c) An example of self-occlusions in ‘Gobangaki.’ (d) 3D model of ‘Ooniwa’ at maturity. (e) Horizontal sections of 3D models. Scores and descriptors in (a) and (b) are based on Fruit Tree Experiment Station of Hiroshima Prefecture (1979).

Maeda et al. (2018) phenotyped persimmon fruit by chronologically applying an elliptic Fourier descriptors-based PCA method for 2D images of 153 *D. kaki* cultivars and two related species. The 2D imaging method successfully represented the ratio of length to diameter and the fruit shape of the distal (apex) end, whereas components representing vertical and horizontal grooves were not detected. In fact, when the entire fruit is photographed from one direction, as in Maeda et al. (2018), the grooves in persimmon fruit are often self-occluded (Fig. 1c); thus, measurement by 2D-based methods is difficult. Ding et al. (2000) developed a 3D morphological analysis method for persimmon fruit. They captured the whole fruit contours using a CCD laser sensor system; however, there were few data points, and development of the unique persimmon fruit characteristics, such as horizontal and vertical grooves, have not yet been fully evaluated.

We previously showed potential applicability of photogrammetry-based 3D phenotyping of groove depths in persimmon fruit; shape measurements by geometric processing of the obtained 3D models represented the genetic diversity in the pattern of vertical groove depth (Kusumi et al., 2022). In this study, we applied a high-resolution 3D scanner to model the full growth of persimmon fruits with diverse shapes and to obtain regulatory information regarding fruit morphological development. In addition, transcriptome analysis was conducted to characterize the molecular mechanism of vertical groove regulation.

## Materials and Methods

### Plant materials

Fruit shapes were assessed for 51 *D. kaki* cultivars planted in the Kyoto Farmstead of the Experimental Farm of Kyoto University (Kyoto, Japan) (Table 1). Four to six fruits were sampled from each cultivar every 2 weeks, starting from the flowering stage in mid-May until the end of growth stage 1 in mid-August, and every 4 weeks after that until maturity, which varied among cultivars from the end of September to the beginning of November. The sampling interval was adjusted because it is known that the persimmon fruit growth follows a double-sigmoid curve, with growth stage 1 corresponding to the period when fruit development is drastically driven by active cell division and cell expansion (Nii, 1980), and its shape does not change rapidly after approximately 10 weeks after blooming (WAB). However, due to the rottenness during the storage, ‘Mizushimagosho’ of 16 WAB contained only 2 fruits.

**Table 1.** Cultivars used in this study.

| No. | Cultivar | Vertical groove | Horizontal groove | Determining timing (WAB) |  |  |  |  |
| --- | --- | --- | --- | --- | --- | --- | --- | --- |
|  |  |  |  | Gl#20-89 | Ro#20-89 | L/W | Horizontal groove position | Horizontal groove depth |
| 1 | Zenjimaru |  |  | 2 | 2 | 12 |  |  |
| 2 | Hazegosho |  |  | 2 | 6 | 8 |  |  |
| 3 | Maekawajiro | * |  | 2 | 2 | 4 |  |  |
| 4 | Kurogaki |  |  | 4 | 4 | 4 |  |  |
| 5 | Aburatsubo |  |  | 4 | 8 | at maturity |  |  |
| 6 | Amayotsumizo | * |  | 6 | 4 | 8 |  |  |
| 7 | Aizumishirazu |  |  | 6 | 6 | 8 |  |  |
| 8 | Fujiwaragosho |  |  | 4 | 4 | 4 |  |  |
| 9 | Kikuhira |  | * | 4 | 0 | 4 | 4 |  |
| 10 | Koyowase |  |  | 4 | 6 | 4 |  |  |
| 11 | Nagara |  | * | 2 | 8 | at maturity | 12 |  |
| 12 | Matsumotowasefuyuu |  |  | 2 | 4 | 2 |  |  |
| 13 | Totsutanenashi |  |  | 6 | 6 | 6 |  |  |
| 14 | Shinshuu |  |  | 2 | 6 | 20 |  |  |
| 15 | Sakata | * | * | 0 | 0 | 8 | 12 |  |
| 16 | Tonewase |  |  | 4 | 0 | 6 |  |  |
| 17 | Atago |  |  | 4 | 6 | 4 |  |  |
| 18 | Gionbou | * |  | 6 | 4 | 8 |  |  |
| 19 | Kiyosu-tanenashi |  |  | 4 | 10 | 6 |  |  |
| 20 | Ooyotsumizo | * |  | 6 | 4 | 4 |  |  |
| 21 | Jisha |  |  | 6 | 4 | 8 |  |  |
| 22 | Ichidagaki |  |  | 2 | 8 | 8 |  |  |
| 23 | Saijou | * |  | 8 | 10 | at maturity |  |  |
| 24 | Dojohachiya |  |  | 6 | 0 | 6 |  |  |
| 25 | Ootanenashi |  |  | 4 | 0 | 4 |  |  |
| 26 | Yokono |  |  | 2 | 4 | 8 |  |  |
| 27 | Fuji |  |  | 4 | 4 | 10 |  |  |
| 28 | Jirou | * |  | 4 | 4 | 0 |  |  |
| 29 | Yamato |  |  | 6 | 10 | 12 |  |  |
| 30 | Fudegaki |  | * | 4 | 4 | at maturity | 10 |  |
| 31 | Komino |  |  | 4 | 6 | 4 |  |  |
| 32 | Amahyakume |  |  | 6 | 16 | 16 |  |  |
| 33 | Okugosho |  |  | 4 | 6 | 10 |  |  |
| 34 | Mino |  |  | 2 | 6 | 10 |  |  |
| 35 | Yoshino |  |  | 4 | 4 | 4 |  |  |
| 36 | Yotsumizo | * |  | 8 | 2 | 2 |  |  |
| 37 | Tenpougai | * | * | 2 | 0 | 4 | 10 | 16 |
| 38 | Hiratanenashi |  |  | 6 | 2 | 4 |  |  |
| 39 | Tamopan |  | * | 4 | 0 | 0 | at maturity | 16 |
| 40 | Fuyuu |  |  | 4 | 8 | 4 |  |  |
| 41 | Iwasedo | * |  | 6 | 0 | 4 |  |  |
| 42 | Midai | * | * | 0 | 4 | 4 | 10 |  |
| 43 | Hagakushi |  |  | 2 | 10 | 10 |  |  |
| 44 | Sangokuichi |  |  | 2 | 0 | 8 |  |  |
| 45 | Kuramitsu |  |  | 2 | 0 | 10 |  |  |
| 46 | Mizugaki/suishi |  |  | 6 | 0 | 8 |  |  |
| 47 | Ooniwa | * |  | 0 | 0 | 8 |  |  |
| 48 | Kawabata | * |  | 0 | 0 | 20 |  |  |
| 49 | Keizanbanshi | * |  | 2 | 4 | 2 |  |  |
| 50 | Mikado | * |  | 8 | 4 | 4 |  |  |
| 51 | Mizushimagosho | * | * | 8 | 8 | 4 | 8 |  |

Asterisks indicates the presence of vertical or horizontal groove.

### Acquiring 3D models

A 3D Scanner Coordinate Measuring Machine (VL-500; KEYENCE, Osaka, Japan) was used to develop 3D models of the persimmon fruit. Fruit was sampled with the peduncle at the first 2 sampling periods, which were 0 and 2 WAB, and the fruit was scanned with the peduncle. After 4 WAB, the fruit calyx was removed to scan the precise features around it. Each persimmon fruit was set on a rotating table, and 3D images were obtained every 30 or 60 degrees; the interval at which images were taken (30 or 60 degrees) was based on sample complexity. The turning angle was optimized according to the fruit shape complexity, and image acquisition was iterated until a high-quality 3D model was obtained. This step was done for both side of each fruit, i.e., the proximal end and distal end, which led to an intact fruit 3D model when they were fused (Fig. 1c). The final model was saved as an STL file. All 3D models developed in this study are available at https://doi.org/10.6084/m9.figshare.22001908.

### Image processing

MeshLab version 2021.05 (Cignoni et al., 2008), trimesh version 3.17.1 (https://github.com/mikedh/trimesh), and OpenCV version 3.1.0 (Bradski, 2000) were used to process 3D models and measure shape features, and the code is available at https://github.com/pomology-ku/persimmon-fruit-3D-measurement. Regarding the samples from 0 and 2 WAB, peduncles were manually removed using MeshLab. For all fruit models, the Z-axis was aligned to the proximal-distal line. Horizontal sections in the XY-plane were obtained by dividing the fruit model into 100 equal parts along the Z-axis. Additionally, 30 vertical sections were acquired by slicing the model by every 6° from the centroid origin.

Based on horizontal sections, vertical groove and roundness parameters were measured (Fig. 2a and Supplementary Fig. 1a). A parameter that represented vertical groove depth, Groove Index (GI), and Roundness (Ro) were calculated by the following equations:

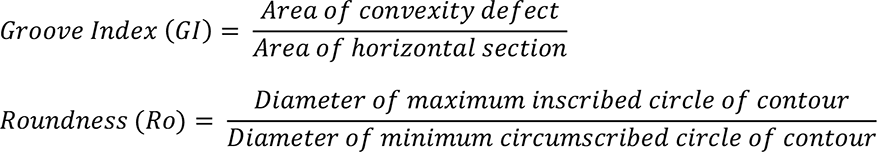

**Figure 2.**
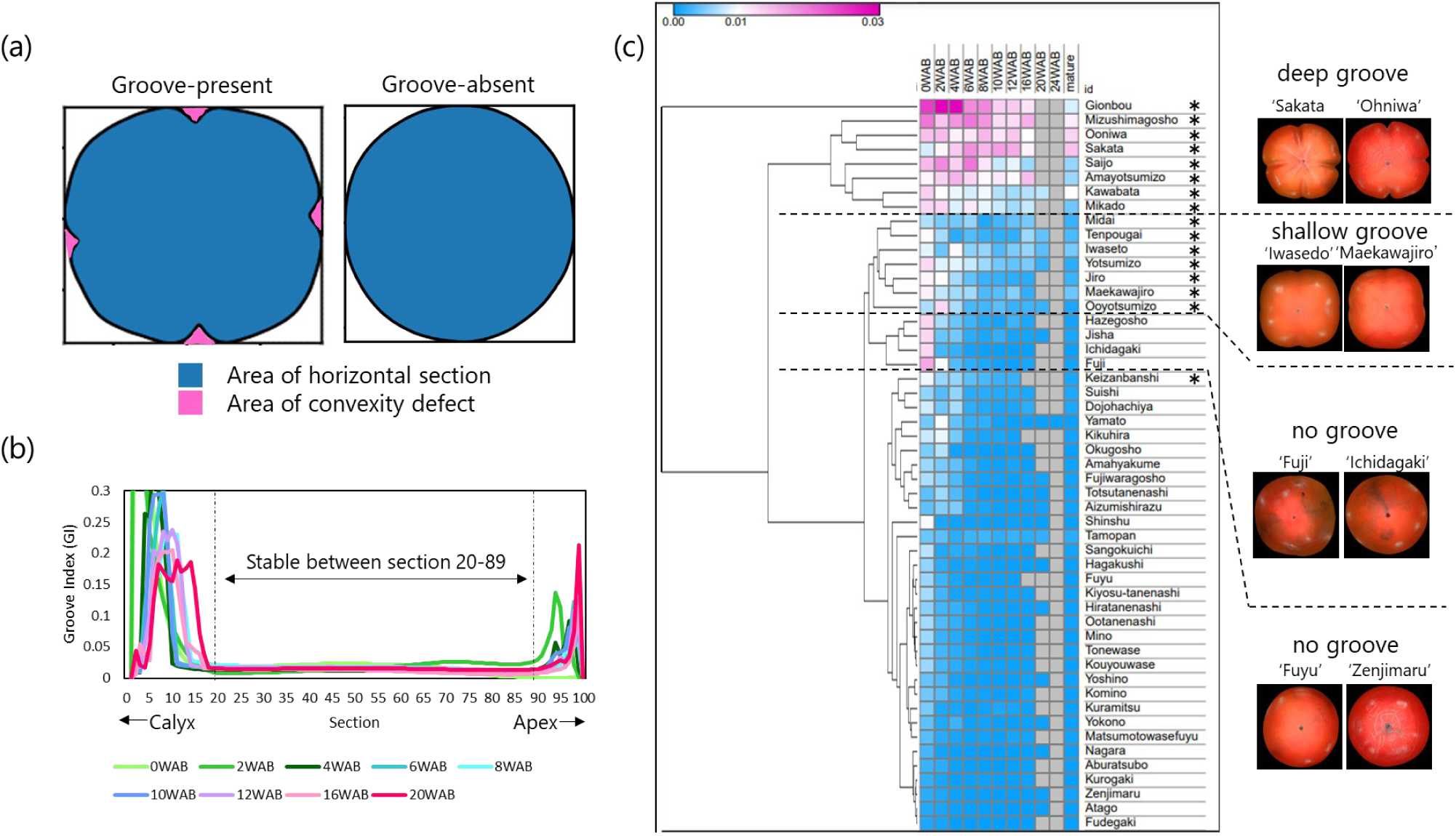
Characterization of vertical groove development. (a) Schematic of the measurement method used to determine groove index (GI). (b) Transition of GI in ‘Ooniwa’ throughout developmental stages. (c) Heatmap of the transition of GI#20-89, with asterisks indicating cultivars classified as having vertical grooves by Fruit Tree Experiment Station of Hiroshima Prefecture (1979) and visually verified when no recorded reference was available.

Length to width (L/W) ratio and horizontal grooves were evaluated from transverse sections. L/W ratio was calculated by the average of L/W from 30 sections (Fig. 3a). After detecting horizontal grooves based on curvature change, groove location was identified as the angle between the proximal–distal line and the line connecting the origin and detected groove. Groove location was averaged across 30 sections for each sample. It should be noted that horizontal groove measurement was conducted for eight cultivars with visible horizontal grooves (Table 1).

**Figure 3.**
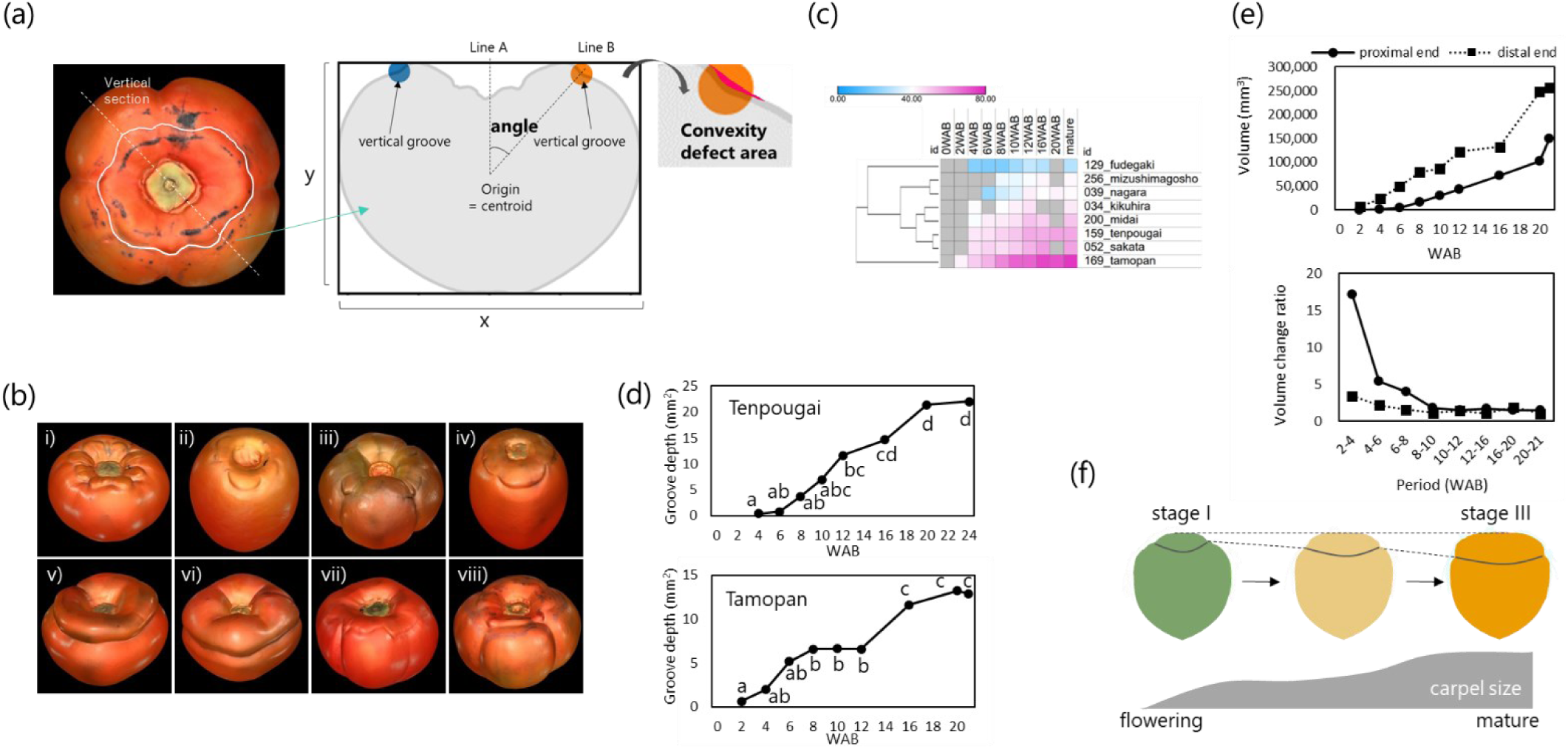
Characterization of horizontal groove development. (a) Schematic of the measurement method used to determine horizontal grooves. (b) 3D models of (i) ‘Kikuhira,’ (ii) ‘Nagara,’ (iii) ‘Sakata,’ (iv) ‘Fudegaki,’ (v) ‘Tenpougai,’ (vi) ‘Tamopan,’ (vii) ‘Midai,’ and (viii) ‘Mizushimagosho.’ (c) Heatmap of the transition of groove position (angle between line A and line B in Figure 3a). (d) Transition patterns of groove depth in ‘Tenpougai’ and ‘Tamopan.’ The black line indicates the horizontal groove depth. (e) Transition of volume and volume increase ratio in ‘Tamopan.’ (f) Proposed developmental model for horizontal grooves.

Finally, Tukey–Kramer’s multiple comparison test (*p* < 0.05) was performed among different developmental stages to determine when these morphological parameters were fixed. The stage when the shape parameter score had no significant difference from mature fruit was considered the stage at which shape attributes were determined.

### Microscopic observation

Five fruits from each of four cultivars (‘Mizuhimagosho,’ ‘Saijou,’ ‘Amayotsumizo,’ and ‘Fuyuu’), which exhibited various GI at blooming, were sampled at anthesis in mid-May 2022. They were then fixed in the fixative FAA (formalin: acetic acid: ethanol: distilled water = 2:1:10:7 v/v/v/v), followed by dehydration in a graded ethanol series. Samples were cryo-embedded in blocks and cut into 6- or 8-µm-thick sections according to Kawamoto (1990). Sections stained with toluidine blue were observed with an optical microscope (BX53FL, OLYMPUS, Tokyo, Japan).

Cell numbers and sizes were calculated as follows: first, baselines connecting the tips of seeds inside a fruit were drawn (Fig. 4d). Second, line G (grooved zone) or N (non-grooved zone) was set from baseline to the point at the contour of the fruit where the groove was deepest, or the point where the distance to the baseline was the longest. Finally, number and size of cells that overlap with line G or N were measured. Parenchyma and tannin cells were independently measured (Fig. 4e). To reduce the effect of outliers on cell size, the upper and lower 5% of data were removed, and mean was calculated with the other 90% of the data. Wilcoxon rank sum test was conducted to assess if there was significant difference between groove-present and -absent zones.

**Figure 4.**
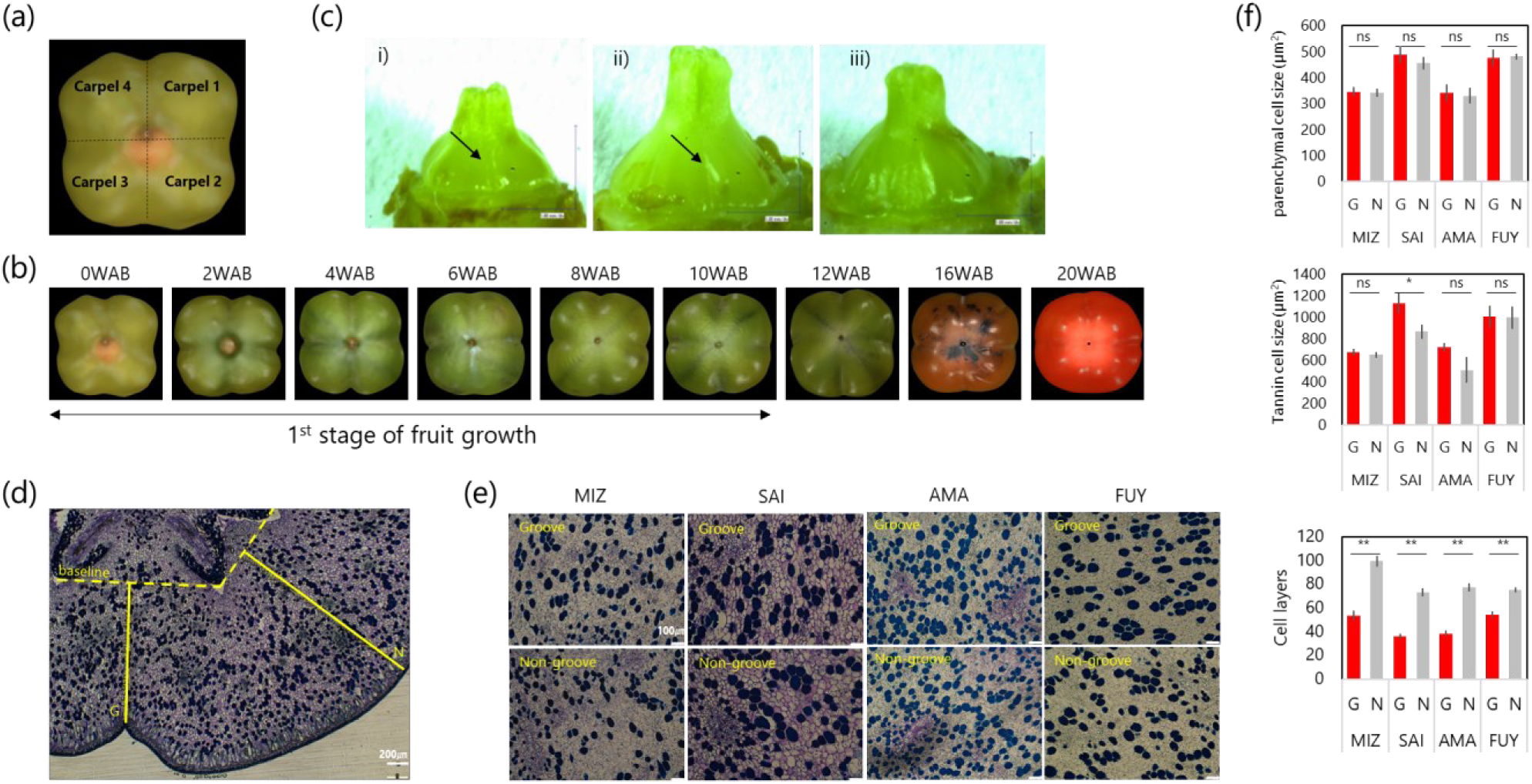
Morphological characterization of groove tissue. (a) 3D model of ‘Gionbou’ at anthesis showing grooves at the boundaries of the fused carpels. (b) Chronological change of groove development in ‘Gionbou.’ (c) Ovaries of (i) ‘Amayotsumizo,’ (ii) ‘Mizushimagosho,’ and (iii) ‘Fuyuu’ at three weeks before anthesis (23 April 2022). Black arrowheads indicated grooving lines. (d) Measurement of cell number and area in the grooved and non-grooved zones using baselines set at the tip of developing seeds. (e) Microscopy analysis performed on grooved and non-grooved zones for four cultivars. MIZ, SAI, AMA, and FUY represent ‘Mizushimagosho,’ ‘Saijou,’ ‘Amayotsumizo,’ and ‘Fuyuu,’ respectively. Blue-dyed cells represent tannin cells. (f) Comparison of parenchyma cell size, tannin cell size, and cell layers measured in each zone (grooved, G, and non-grooved, N) in four cultivars. Single and double asterisks indicate significant differences by paired t-test at p < 0.05 and p < 0.01, respectively.

### Transcriptome analysis

Fruits were sampled from eight cultivars (‘Zenjimaru,’ ‘Fuyuu,’ ‘Amayotsumizo,’ ‘Fujiwaragosho,’ ‘Gionbou,’ ‘Saijou,’ ‘Iwasedo,’ and ‘Mizushimagosho’) at 4 WAB (10 June 2022). Those cultivars were selected because the vertical groove depth plateaus at around or after 4 WAB (except ‘Zenjimaru,’ which reached plateaus at around 2 WAB). The grooved and non-grooved zones for each sample (Fig. 5a) were cut from each fruit, and the peel and developing seeds were removed. Each tissue from each fruit was considered a single biological replicate, and four or five fruits (replicates) were prepared for each cultivar and tissue type.

**Figure 5.**
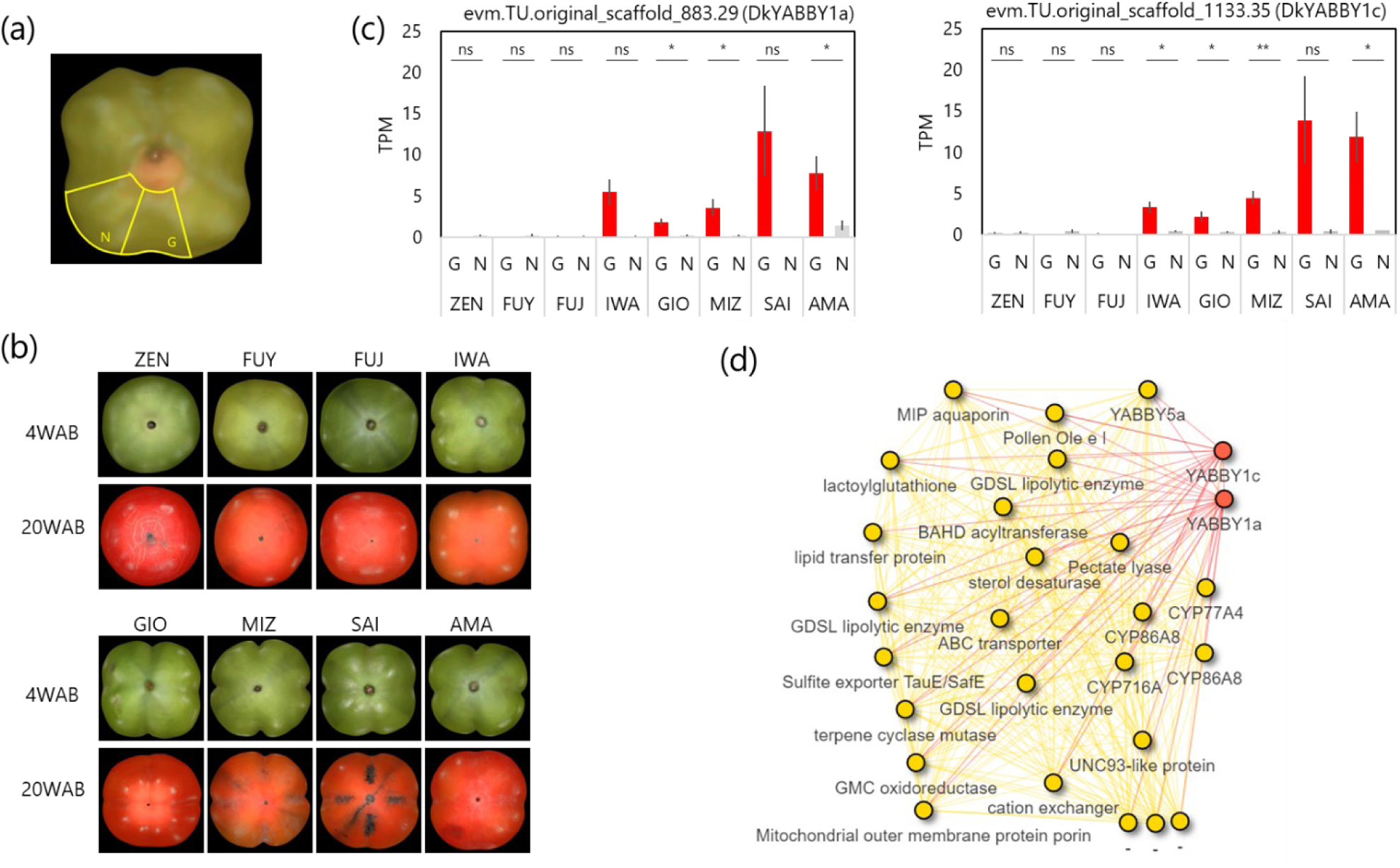
Transcriptome analysis of vertical groove development. (a) Grooved (G) and non-grooved (N) zones from which RNA was extracted. (b) Vertical groove depths at 4 and 20 WAB. (c) Expression patterns of *DkYABBY1a* and *DkYABBY1c*. Single and double asterisks indicate significant differences at p < 0.05 and p < 0.01 by paired t-test, respectively. (d) Gene co-expression network for *DkYABBY1a*/*1c*. The full network of this module is presented in Supplementary Figure S8. ZEN, FUY, FUJ, IWA, GIO, MIZ, SAI, and AMA represent ‘Zenjimaru,’ ‘Fuyuu,’ ‘Fuji,’ ‘Iwasedo,’ ‘Gionbou,’ ‘Mizushimagosho,’ ‘Saijou,’ and ‘Amayotsumizo,’ respectively.

Total RNA was extracted by the hot borate method (Wan and Wilkins, 1994). Extracted RNA was quantified using a Qubit 4 fluorometer (Invitrogen, Carlsbad, CA, USA). In total, 500 ng of total RNA was used to prepare mRNAseq libraries. mRNAseq libraries were prepared using the NEBNext® Poly(A) mRNA Magnetic Isolation Module, NEBNext® Ultra™ II RNA Library Prep Kit for Illumina®, and NEBNext® Multiplex Oligos for Illumina® (New England Biolabs, Ipswich, MA, USA) following the manufacturer’s recommendations. The 78 libraries were split into two groups of 40 and 38 to avoid barcode sequence overlap, and they were sequenced in one lane of DNBSEQ (PE150) for each pool at Azenta Inc, Japan (Tokyo, Japan). All RNA samples used for RNAseq analysis are listed in Supplementary Table 1. All sequence data are available in the DDBJ Sequenced Read Archive (accession number DRA015493).

All sequences were pre-processed using fastp (Chen et al., 2018) with a minimum read length threshold of 35 bp. The reads from each library were aligned to the *Diospyros oleifera* reference genome (Suo et al., 2020) using STAR with default parameters (Dobin, et al., 2013). Alignments were sorted using SAMtools (Li et al., 2009), and the number of reads mapped to each contig was counted from the BAM files using HTseq-count (Putri et al., 2022). Gene expression level was represented as transcripts per million (TPM).

Differential expression between grooved and non-grooved zones in groove-present cultivars was evaluated using Deseq2 version 1.34.0 (Love et al., 2014) with the cultivar factor being input as cofactor of the model. A false discovery rate of 0.05 was set to define differentially expressed genes (DEGs). Genes in the *D. oleifera* reference genome was functionally annotated using eggNOG 5.0 (Huerta-Cepas et al., 2019) with default parameters. Gene ontology enrichment analysis of the groove-associated DEGs was conducted using goseq (Young et al., 2010) with a false discovery rate of 0.1 as the significance threshold. A gene co-expression network was constructed using WGCNA (Langfelder and Horvath, 2008) with a TPM matrix for genes with TPM > 1 and the following parameters: minModuleSize = 30, mergeCutHeight = 0.25. Soft thresholding power was set to 9, with a scale-free topology fitting index of R^2^ > 0.85. The gene module containing the *YABBY* homologs and network edges with topological overlap greater than 0.30 was visualized using R package visNetwork version 2.1.2 (https://github.com/datastorm-open/visNetworkg).

## Results

### Three-dimensional modeling of persimmon fruit

The VL-500 3D scanner developed fruit 3D models that precisely reflected morphological features of persimmon fruit, including horizontal and vertical grooves (Fig. 1c, d). The polygon meshes of fruit 3D models included at least 120 vertices per mm^2^, which is sufficiently fine to evaluate fruit shape at flowering (*ca.* 4.08 mm in longitudinal diameter) in the cultivar with the smallest ovary at flowering, ‘Totsutanenashi.’ Examples of horizontal sections from ‘Ooniwa’ and ‘Saijou’ are shown in Fig. 1d. The slice contours showed clear grooves, and the models included genetic differences in groove depth, including in ‘Ooniwa,’ which has deeper grooves and a deeper cavity at the grooved zone. These general features of 3D models supported the feasibility of evaluating complex shape attributes based on cross sections of the 3D models.

### Characterization of vertical groove disappearance using 3D modeling

To acquire a reliable metric to measure the shape attributes, GI and Ro were visualized over cross sections from the proximal to the distal end (Fig. 2a and Supplementary Fig. 1a). The contours in horizontal sections near the proximal or distal ends were often rough and not stable because of concavity at both ends (Fig. 2b and Supplementary Fig. 1b). Thus, GI and Ro were only evaluated from section #20-89, and were named GI#20-89 and Ro#20-89, respectively.

Fig. 2c represents the chronological change in GI#20-89 from all cultivars as clustered by Euclidean distance. The cultivars with vertical grooves clustered in conventional categories but were largely separated into two groups according to change pattern. We observed that all analyzed cultivars had vertical grooves in fruit at flowering, and GI#20-89 decreased as fruit developed. By 6 WAB at the latest, the groove shape of 47/51 analyzed cultivars showed GI#20-89 values consistent with the value at maturity, with the remaining four cultivars determined at 8 WAB (Table 1). Therefore, vertical groove depth was mainly determined in the initial stage of fruit development, at which time active cell division rather than cell expansion takes place.

### Roundness and its association with vertical groove depth

Roundness was also evaluated from transverse sections. Supplementary Fig. 1 shows a chronological change in Ro#20-89 for all cultivars, as classified by Euclidean distance. Generally, Ro#20-89 tended to increase with development, which indicated that the fruit became round during development. Interestingly, cultivars with vertical grooves generally clustered together; these cultivars tended to be less round. In most cultivars, roundness reached at the maturity level by 10 WAB (Table 1).

To further investigate the relationship between the groove depth and roundness, Spearman’s rank correlation analysis was performed. Through developmental stages, groove depth and roundness were significantly correlated with each other (*p* < 0.01) (Supplementary Fig. 2a), which indicated that grooved fruit tends to be square-shaped. At 0 WAB, the correlation coefficient was −0.518. Then, the coefficient gradually increased and reached −0.743 at maturity (Supplementary Fig. 2b). This result implies a common mechanism underlying both vertical groove depth and roundness.

### Horizontal groove and L/W development during fruit growth

Horizontal grooves and L/W development were evaluated using 30 vertical sections of each fruit (Fig. 3a). Supplementary Fig. 3a shows a chronological change in L/W for all cultivars, as classified by Euclidean distance. L/W remained low through the development of flat cultivars; however, it increased toward maturity in oblong cultivars. ‘Fudegaki’ and ‘Nagara,’ which have oblong fruit, exhibited drastic changes in L/W. L/W was determined within the first developmental stage in most cultivars, but there were several cultivars whose L/W was not determined by 24 WAB. Cultivars with late determination included many oblong cultivars, such as ‘Nagara,’ ‘Fudegaki,’ ‘Saijou,’ and ‘Aburatsubo.’

Detecting horizontal grooves was difficult and produced many missing values. This is because the grooves typically did not cover the entire fruit, or the curvature change at the grooves in vertical sections were not as large as other depressions in fruits (Fig. 3b). Additionally, horizontal grooves near the proximal end might be missing from the samples harvested at 0 and 2 WAB because those samples had peduncles when scanned. Fig. 3c shows the chronological change in horizontal groove point angles, which represented the location of horizontal grooves, for fruit from eight cultivars, as classified by Euclidean distance. Generally, the score increased as the fruit grew, which indicated that horizontal groove positions tended to move from the proximal to the distal end. In particular, ‘Tamopan,’ which has an “extremely big” horizontal groove (Fruit Tree Experiment Station of Hiroshima Prefecture, 1979), had a distinctive pattern of groove development; the groove position kept changing toward the distal end throughout fruit development. However, ‘Kikuhira’ and ‘Mizushimagosho’ did not show any significant differences in groove position since horizontal groove emergence. Groove development became stable at 10 WAB in ‘Tenpougai,’ ‘Midai,’ ‘Nagara,’ and ‘Fudegaki,’ and at 12 WAB in ‘Sakata’ (Supplementary Fig. 4).

Additionally, changes in horizontal groove depth were also evaluated in ‘Tenpougai’ and ‘Tamopan.’ The depth continued to increase from anthesis, but the increase suspended at mid-growth and showed double-sigmoid curve patterns (Fig. 3d). Fruit volume analysis in ‘Tamopan’ showed that the distal portion, which was beneath the groove, exhibited a double sigmoid-like curve, but the proximal portion, which was above the groove, did not follow this pattern and continued to develop (Fig. 3e). Besides, the volume increase ratio was larger in the proximal portion than in the distal portion throughout the developmental stages except at 20 WAB. Especially in the early growing period, the differences were much larger (Fig. 3e).

### Developmental analysis of vertical groove variation

In this study, we conducted an in-depth analysis of the genetic diversity of vertical grooves that many cultivars possess. Persimmon generally have four carpels, and vertical grooves develop at carpel fusion zones (Fig. 4a). Most persimmon cultivars show various degrees of a vertical groove at anthesis, regardless of vertical groove presence or absence at maturity (Fig. 2c). However, the groove depth can become shallower during development depending on genotype (Fig. 4b).

To test if the groove depth is primarily determined by cell division or cell expansion, groove development for ovarian tissues was characterized before and at anthesis in four cultivars with various groove depths. At approximately 3 weeks before anthesis, groove lines were visible along the carpel fusion lines in ‘Amayotsumizo’ and ‘Mizushimagosho,’ but not in ‘Fuyuu’ (Fig. 4c). Microscopic analysis revealed no significant difference between cell sizes in groove-present and -absent zones regardless of the cell types and cultivars except tannin cell size in ‘Saijou’. However, there was a significant increase in the whole cell numbers counted for both tannin cells and parenchyma cells in groove-absent tissue compared with groove-present tissue in all cultivars (Fig. 4-f; p < 0.01 in the paired Student’s *t*-test). The significant difference in parenchyma cell number between groove-present and - absence was observed in all four cultivars (Supplementary Fig. 5; p < 0.01 in the paired Student’s *t*-test). In addition, a significant difference in tannin cell number was also observed in ‘Saijou’ and ‘Amayotsumizo’ (Supplementary Fig. 5; p < 0.05 in the paired Student’s *t*-test). These results indicated that vertical groove formation at anthesis is driven by the variations in cell layers of fruit flesh, mainly of parenchyma tissues, rather than differences in cell size.

We further characterized molecular genetic regulation of vertical groove development using a transcriptomic approach. In differential expression analysis, genetic variation was categorically incorporated into the multi-factor model to standardize differences in the genetic background among cultivars. The differential expression analysis identified 1,883 DEGs between grooved and non-grooved zones in groove-present cultivars. DEGs were typically enriched for functions in cell wall development and cell fate determination (Table 2). To search for candidate regulators, we identified the lowest q-values, which yielded genes with typical functions in fruit development, such as cell wall-related genes and well-characterized transcription factors (Table 3). Specifically, *SVP* and *YABBY* homologs were among the top 20 DEGs with the lowest q-values (Table 3).

**Table 2.**
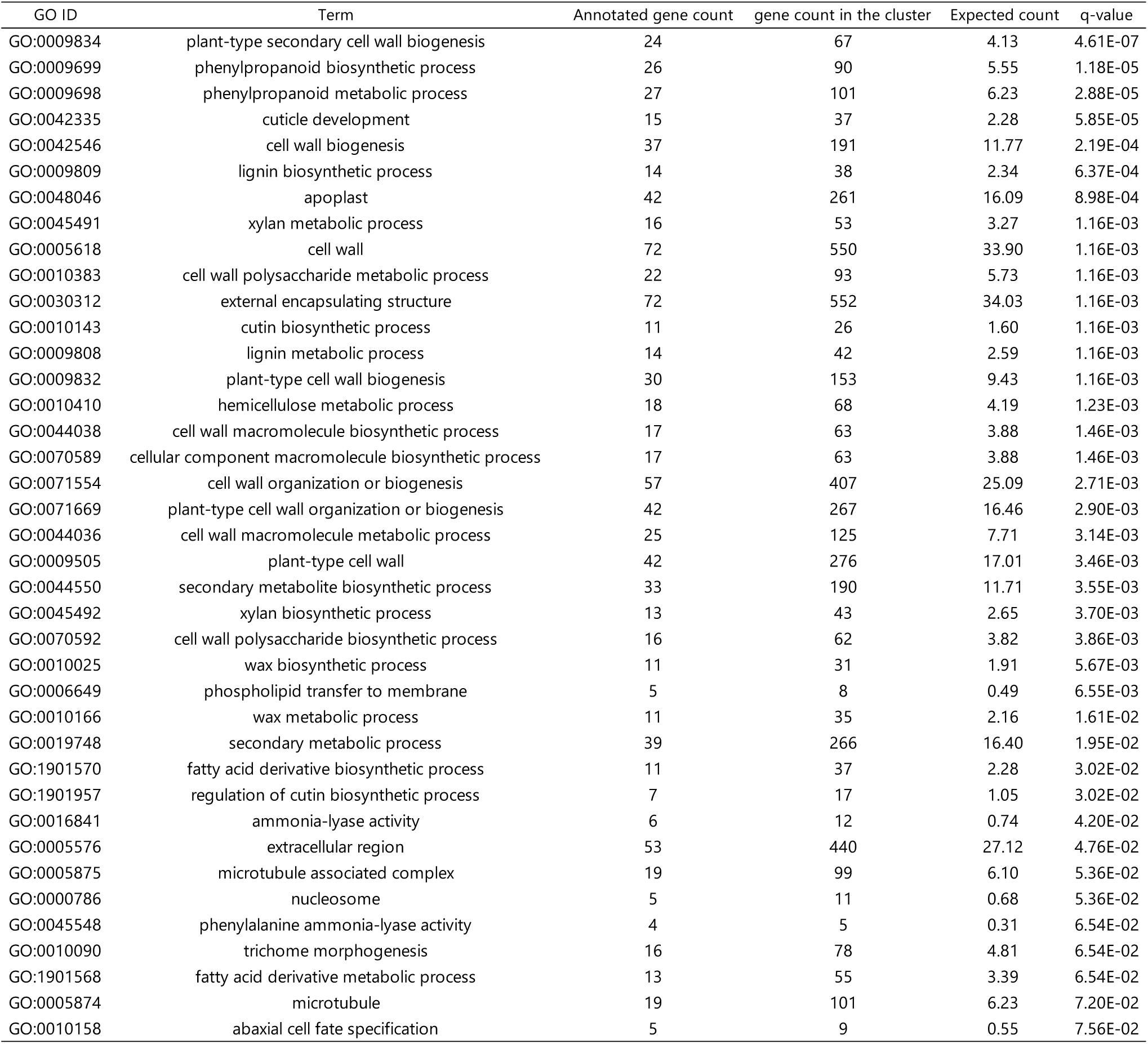
GO enrichment analysis for DEGs.

**Table 3.**
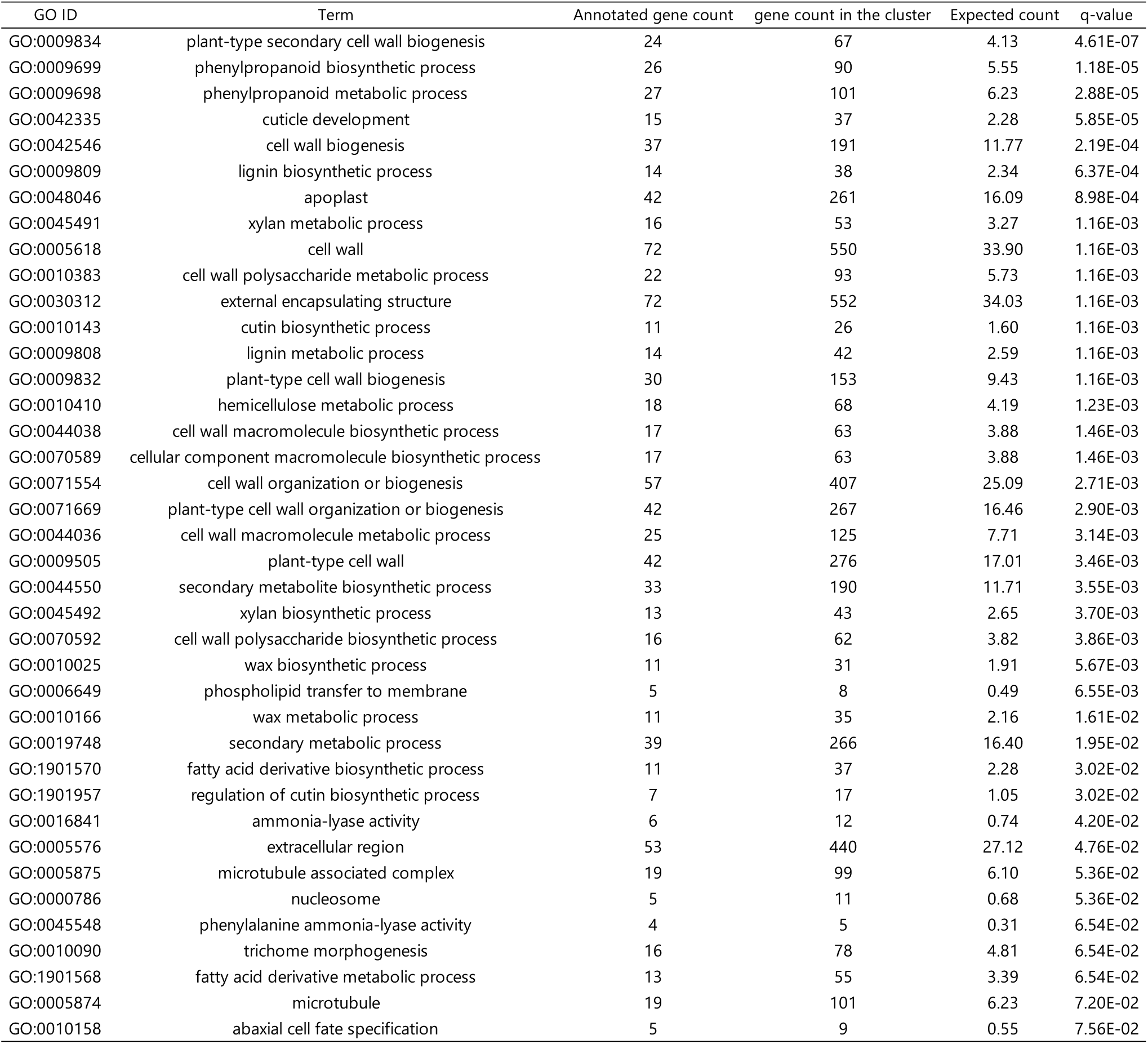
Top 20 genes based on the log2 fold change between grooved and non-grooved zones.

*SVP* is a MADS-box factor that determines floral meristem identity (Gregis et al., 2008). The diploid *Diospyros* reference genome has two *SVP* homologs (Supplementary Fig. 6). One *SVP* homolog (evm.model.original_scaffold_495.37), *DkSVP1*, was the DEG with the second lowest q-value among all genes and showed lower expression in groove than non-groove tissue in groove-present cultivars (Supplementary Fig. 6). The same expression pattern was found in groove-absent cultivars. Interestingly, the other *SVP* homolog, *DkSVP2* (evm.model.original_scaffold_1207.202), generally showed opposite expression compared with *DkSVP1*: there was higher expression in groove than in non-groove tissues.

The *YABBY* gene family is a central regulator of carpel development. In *Arabidopsis*, *YAB1* and *YAB3* are known to have roles in carpel fusion development (Nole-Wilson and Krizek, 2006). The DEGs highly associated with groove development, evm.model.original_scaffold_883.29 and evm.model.original_scaffold_1133.35 (*DkYABBY1a* and *DkYABBY1c*, respectively), were found in the YAB1/YAB3 clade (Supplementary Fig. 7). *DkYABBY1a* and *DkYABBY1c* expression patterns matched the patterns of groove development and genetic variation. They were significantly more expressed in the grooved zone in most groove-present cultivars, and the mean expression level was higher in cultivars with deeper vertical grooves (‘Saijou’ and ‘Amayotsumizo’); however, there were no significant differences between tissues in the groove-absent cultivars (Fig 5c). Similar expression patterns were also observed in evm.model.fragScaff_scaffold_45.107 (*DkYABBY5a*) and evm.model.original_scaffold_1639.203.1 (*DkYABBY1b*) (Supplementary Fig. 7).

The functions and expression patterns of *YABBY* family genes explained the groove depth variation. Therefore, we further characterized the associated gene networks (Fig. 5d and Supplementary Fig. 8). Genes with first-degree interactions with *YABBY* genes did not include transcription factors. However, many transporter genes and enzyme-coding genes were found, such as *CYP77A4*, which functions in auxin-mediated patterning (Kawade et al., 2018), and *CYP86A8*, which functions in omega-hydroxylation of various fatty acids during development (Wellesen et al., 2008). Collectively, these results identify the *YABBY* family as a potential molecular factor that directly regulates vertical groove development.

## Discussion

### Utilizing 3D scan for geometric phenotyping of fruit

This study showed that 3D models developed by a 3D scanner captured detailed morphological features of persimmon fruit. In particular, fine characteristics of complex features, such as horizontal and vertical grooves, were clearly reproduced (Fig. 1c and 3b). Conventional persimmon fruit phenotyping was previously conducted based on 2D imaging (Maeda et al., 2018, 2019; Kusumi et al., 2022), which did not capture all shape attribute information. In this study, we obtained morphological information along various axes. For example, vertical grooves are more prominent near the proximal or the distal end in some cultivars, but they are more noticeable in the middle portion of the fruit in other cultivars. However, 2D imaging cannot capture grooves that form near the distal end because it is obscured when imaged from one side of the fruit. In addition, the horizontal grooves do not typically cover the entire fruit circumference, which results in loss of information when conducting 2D imaging. To address these issues, we used a 3D modeling approach that allowed us to integrate information from multiple sections along the proximal– distal line.

### 3D modeling clarifies complex shape development in persimmon fruit

In this study, we observed that the vertical groove depth decreased during growth stage 1, with variations depending on genotype. However, all cultivars had vertical grooves at anthesis, but with differing depths (Fig. 2). Therefore, it was suggested that the formation of vertical grooves occurred during carpel fusion, and the groove pattern at maturity is shaped by combination of the fusion pattern and cell growth patterns after formation, which could be mainly driven by cell divisions.

On the other hand, the results indicated that horizontal groove development in persimmon fruit is promoted by the different growth speeds between fruit portions above and beneath the groove. We observed that the horizontal groove position shifted toward the distal end and the groove continued to deepen throughout fruit development, which could be attributed to vertical growth and horizontal growth, respectively (Fig. 3c and 3d). This demonstrates that both vertical and horizontal development speeds of proximal tissues above the groove were much faster than those of the distal tissue during early fruit growth, and became similar around growth stages 2-3 depending on cultivars (Fig. 3e and 3f). These observations are consistent with the findings of previous studies, which reported that growth at the proximal end in persimmon fruit is more vigorous during stage 1, which could continue until early-mid August (Fujimura, 1935). Collectively, we showed that differences in the horizontal and vertical growth speeds between portions of the fruit, divided by the horizontal groove, explained horizontal groove development and movement of its position (Fig. 3f).

Maeda et al. (2018) reported that fruit L/W ratio was the main contributor to shape variation in longitudinal fruit sections based on PCA of diverse persimmon cultivars. They found that the correlation coefficients between PC1, which represented L/W ratio variability, at maturity and that of several different developmental stages reached 0.9 at the end of June. In this experiment, correlation analysis was conducted between the L/W ratio at maturity and that of different developmental stages. The correlation coefficient reached at 0.9 between 4–6 WAB (Supplementary Fig. 3b). This result was also consistent with that of Sugiura et al. (1977), in which ethanol treatment of persimmon fruits after flowering until late July influenced the fruit L/W ratio. In this study, we further showed that each genotype had variable transitions; the period for determining L/W ratio varied from 0 WAB to maturity depending on cultivar (Table 1). In addition, we found that the L/W ratio was determined later in cultivars with oblong fruit, such as ‘Fudegaki’ or ‘Nagara.’ This probably means that vertical cell division or expansion receive more resources in oblong cultivars, which causes slower transverse development, but horizontal and vertical development can become equal earlier in other cultivars.

### Cell identity control is the most likely mechanism underlying vertical groove development variation

Vertical grooves in fruit are known to emerge in many species, including *Cucumis* and *Prunus* (Paris, 2016; Yousif et al., 2011). Multiple genetic studies have explored the genes that contribute to groove formation, but the molecular mechanism of their development is not fully understood. Yao et al. (2018) found that overexpression of *Malus domestica PISTILLATA* causes vertical groove development on fruit from 12 days after pollination. Du et al. (2022) identified a 1.07-kb deletion of a chromosome segment in groove lines of *Cucumis melo* and proposed that *CmMADS-Box* may be a candidate gene regulating the groove formation. It was also reported that certain *Arabidopsis* mutants with defects in the *YABBY* and *SVP* genes produced flowers with abnormal carpel fusions, which can possibly lead to groove-like structures (Nole-Wilson and Krizek, 2006; Gregis et al., 2009). However, there has been no report to date of *YABBY* or *SVP* directly regulating groove formation in species with fleshy fruit.

Our transcriptome analysis indicated that DEGs between grooved and non-grooved zones were related to cell wall development or cell fate, which is consistent with the observation that grooves were formed by fewer cell layers at carpel margins (Fig. 4). Interestingly, of the DEGs, *SVP* and *YABBY* gene families had low q-values (Table 3). *SVP* is known to be involved in determination of floral organ identity in *Arabidopsis* and tomato (Liu et al., 2007; Thouet et al., 2012). However, *YABBY* controls establishment of abaxial–adaxial polarity in lateral organs (van der Knapp et al., 2014). *YABBY* is a plant-specific transcription factor with several family members (Bowman, 2000); *FIL* and *YAB3* define the abaxial cell fate in lateral organs in *Arabidopsis* (Siegfried et al., 1999). Besides, *YAB2* and *YAB5*, which are expressed specifically in the abaxial side, can also contribute to the polarity formation with less magnificent effect (Stahle et al., 2009; Sarojam et al., 2010). Importantly, *DkSVP1* expression was also high in groove-absent cultivars, whereas *DkYABBY1a* and *DkYABBY1c* expression completely matched the groove patterns, including presence/absence among different cultivars (Fig 5c). These results strongly suggest that *YABBY* homologs are direct regulators of vertical groove development.

The gene co-expression network of *DkYABBY1a/1c* did not include any transcription factors, which indicated that *YABBY*s are upstream regulators of the network (Fig. 5d). Most genes associated with *YABBY*s are transporters and enzymes, and they seemed to be functionally related to cell development. In particular, enzymes related to hydrocarbon metabolism were highly enriched, such as cytochrome P450 proteins (Fukushima et al., 2011) and lipolytic enzymes. *CYP77A4* was reported to be involved in establishing polarity in *A. thaliana* embryos by regulating auxin distribution pattern; thus, it plays an important role in leaf morphogenesis (Kawade et al., 2018). Additionally, pectate lyase was found in the network, which helps cell wall remodeling (Uluisik and Seymour, 2020). Considering the expression pattern, biological function in model species, and co-expression network, it is very likely that the *YABBY* homologs directly regulate vertical groove development. It should be noted that multiple *YABBY* homologs were expressed in a highly correlated manner, which indicates an additional layer of expression regulation underlying the genetic variation in vertical groove development. Further functional analysis, including examination of characteristic paralogs, in persimmon may offer new insights into this process.

## Acknowledgments

We thank Mallory Eckstut, PhD, from Edanz (https://jp.edanz.com/ac) for editing a draft of this manuscript. This work was supported by the Japan Society for the Promotion of Science KAKENHI Grant Number 20K20454 to SN and RT, and 21KK0269 to SN.

