## Supplementary Materials for "Three-dimensional fruit growth analysis clarifies developmental mechanisms underlying complex shape diversity in persimmon fruit"

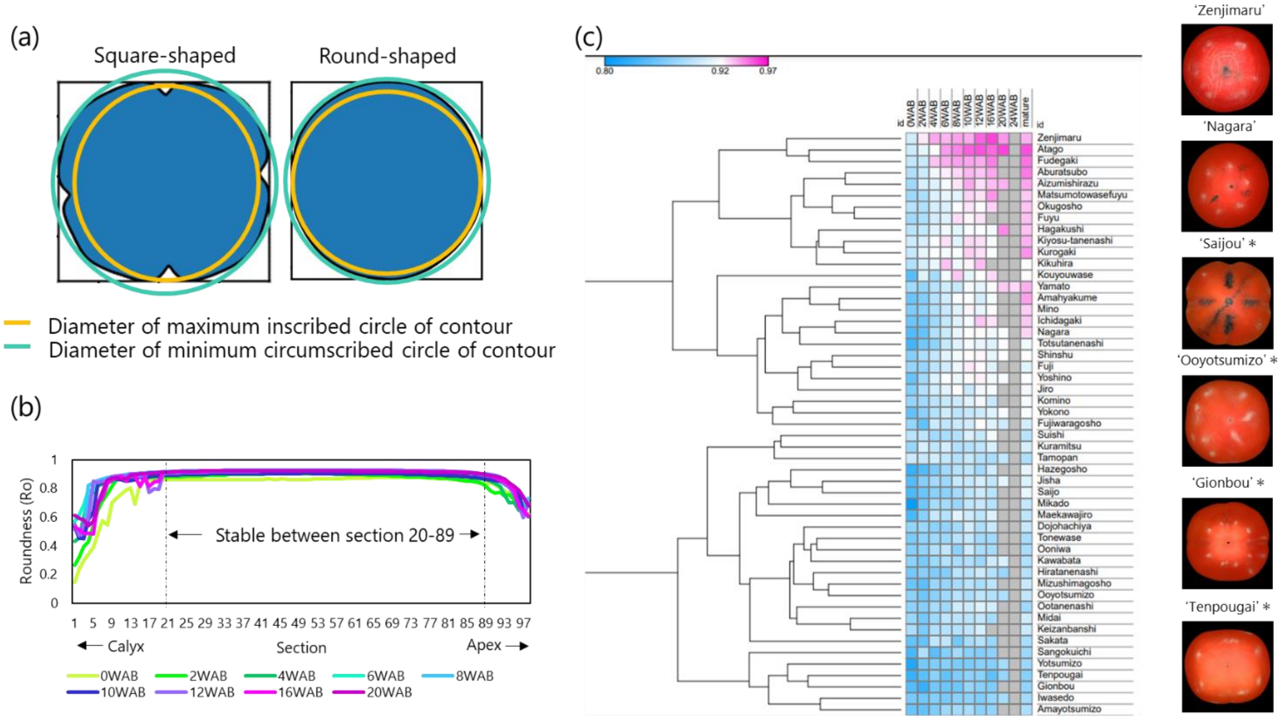


**Supplementary Figure 1. Characterization of roundness development.** (a) Measurement method of Ro. (b) Transition of Ro in 'Ooniwa' from different developmental stages. (c) Heatmap of the transition of Ro#20-89. Asterisks indicate the cultivars classified as having vertical grooves by Fruit Tree Experiment Station of Hiroshima Prefecture (1979) and visually verified when no records were available.

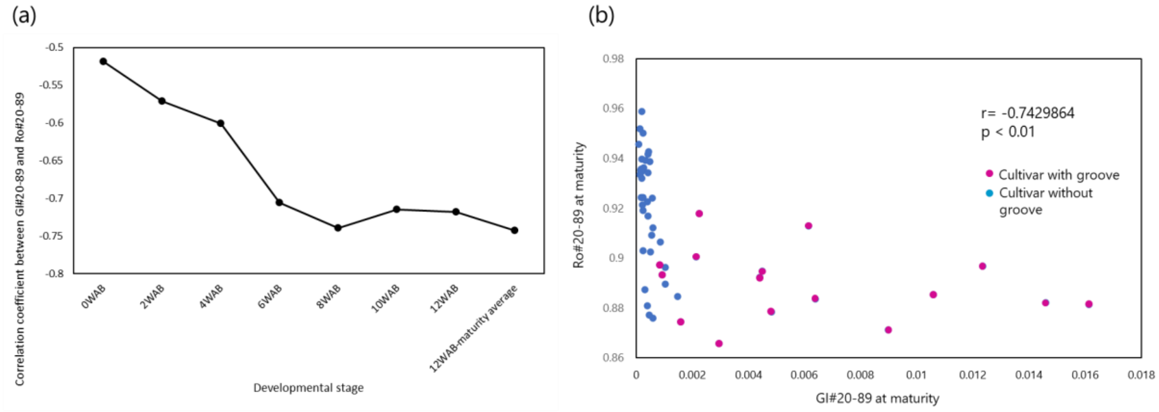


**Supplementary Figure 2. Relationship between groove depth (GI#20-89) and roundness (Ro#20-89).** Relationship (a) through developmental stages and (b) at maturity.

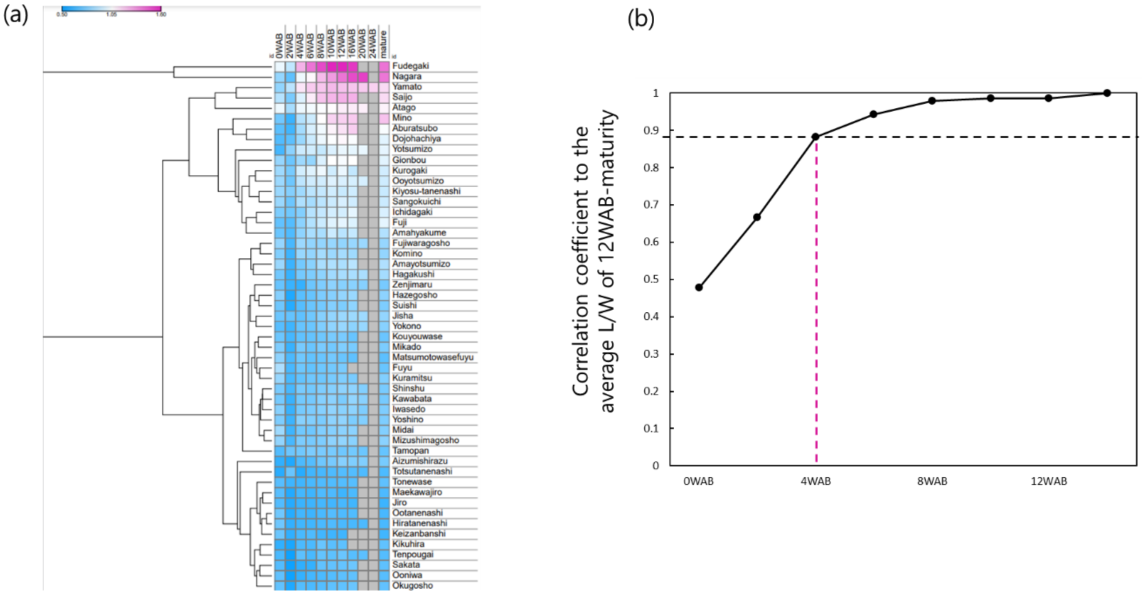


**Supplementary Figure 3. Length/width (L/W) relationships for each cultivar over time. (a)** Heatmap of the transitions of L/W, and (b) correlation coefficients calculated between L/W at each developmental stage and the average L/W from 12 WAB until maturity.

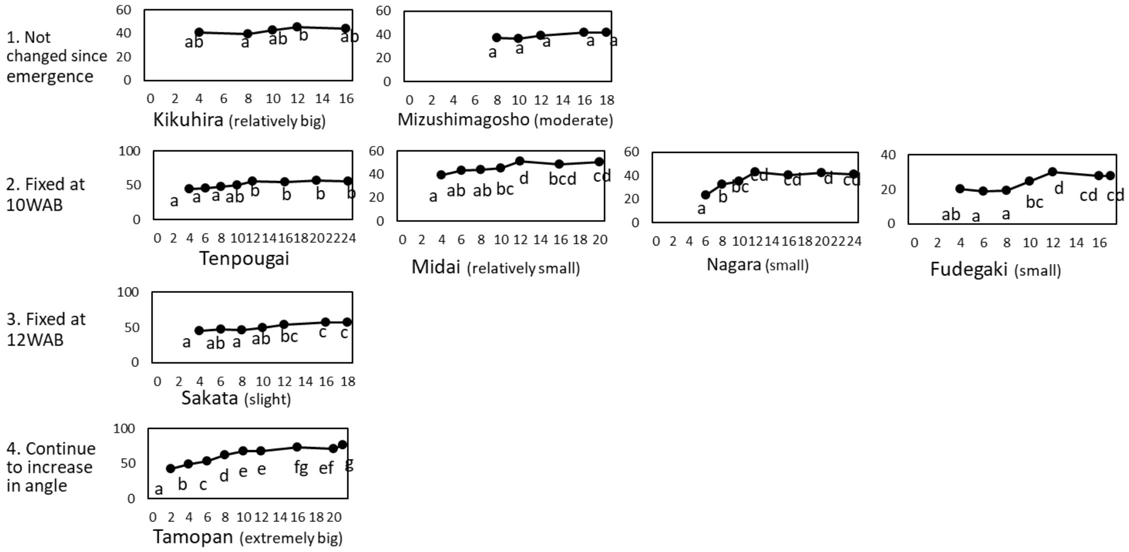


**Supplementary Figure 4. Transition patterns of groove position.**

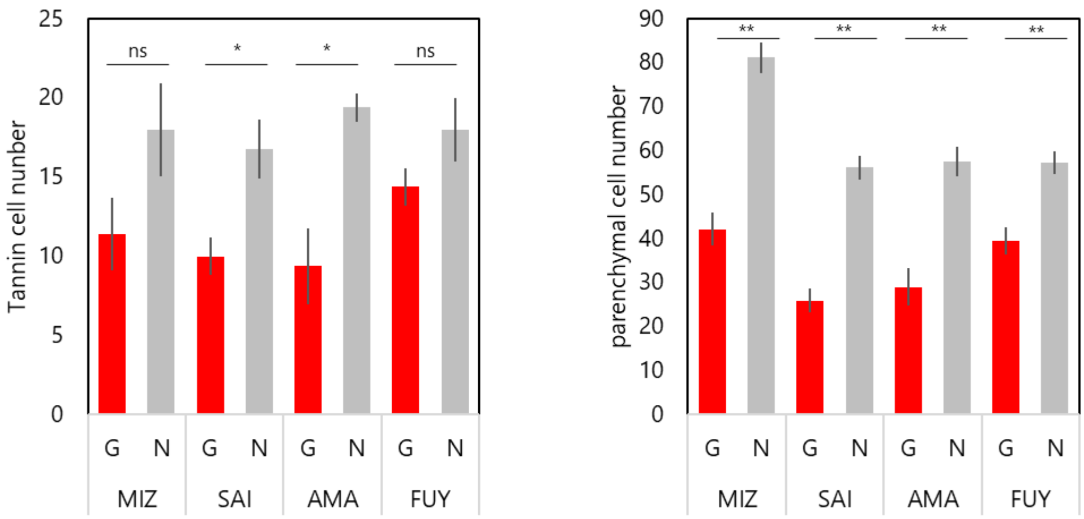


**Supplementary Figure 5. Tannin cell number and parenchymal cell number in grooved (G) and non-grooved (N) zones.**

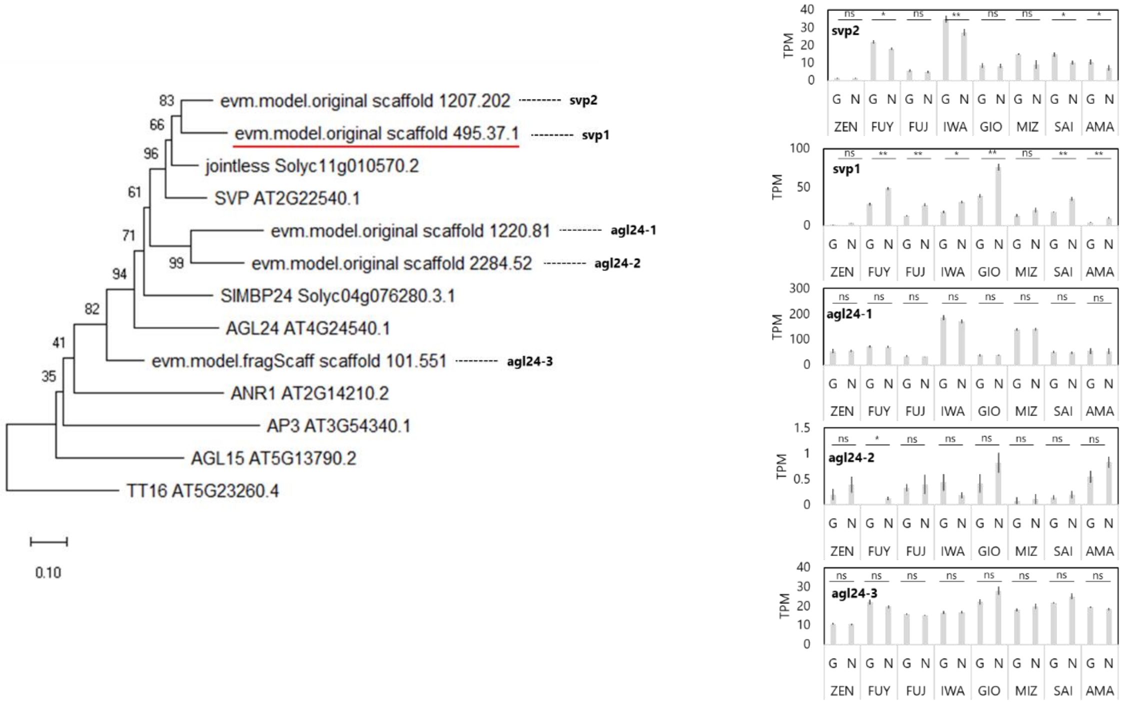


**Supplementary Figure 6. Phylogenetic analysis of identified *SVP* family genes and their expression levels.**

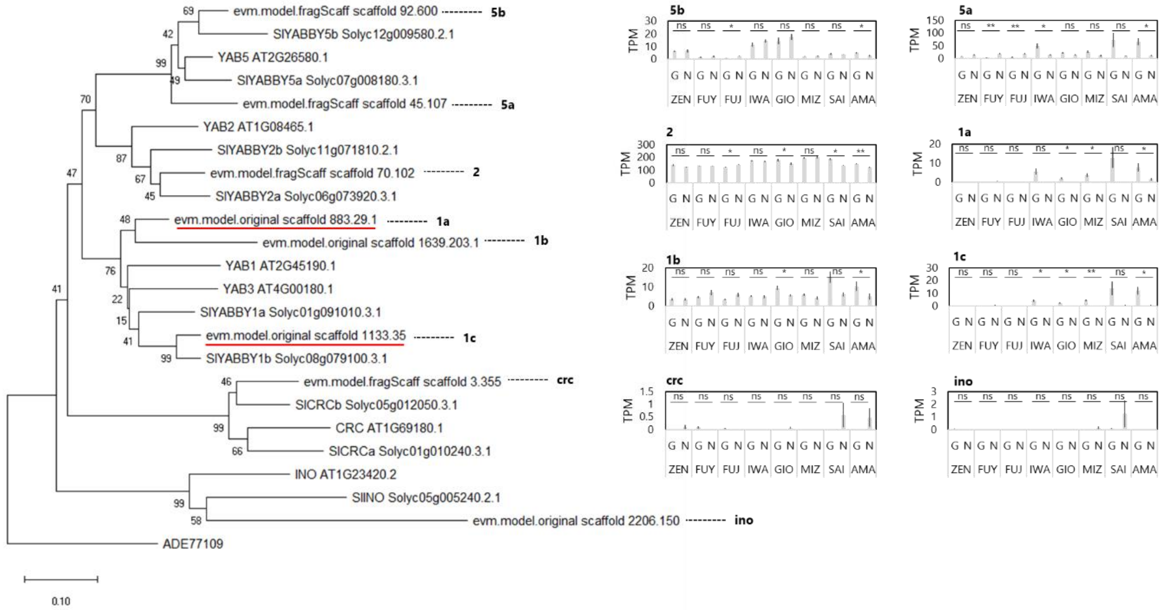


**Supplementary Figure 7. Phylogenetic analysis of identified *YABBY* family genes and their expression levels.**

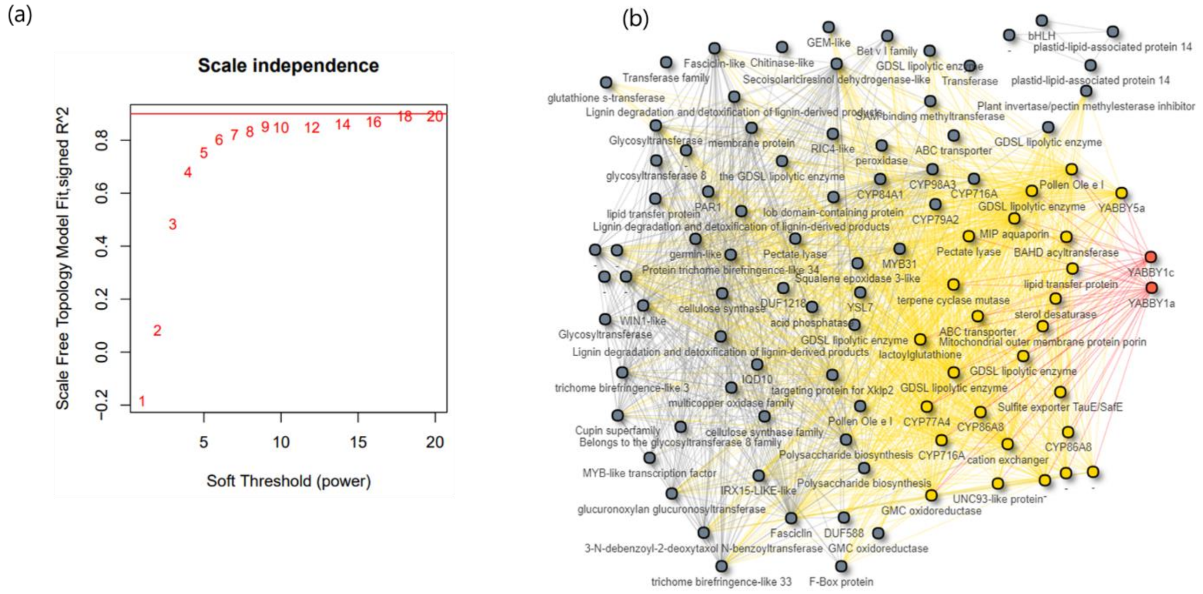


**Supplementary Figure 8. Gene networks associated with *DkYABBY1a* and *DkYABBY1c*.** (a) Scale-free fitting index. (b) Visualization of the full network of modules associated with *DkYABBY1a* and *DkYABBY1c*.

### Supplementary table 1. Summary of RNA-seq libraries constructed in this study

| library name | Run ID | cultivar | tissue | weeks after flowering | replicates ID | number of reads after filtering |
|---|---|---|---|---|---|---|
| 001_G1 | DRR432197 | Zenjimaru | G | 4 | 1 | 16,311,094 |
| 001_G2 | DRR432198 | Zenjimaru | G | 4 | 2 | 16,028,111 |
| 001_G3 | DRR432199 | Zenjimaru | G | 4 | 3 | 15,647,909 |
| 001_G4 | DRR432200 | Zenjimaru | G | 4 | 4 | 14,790,558 |
| 001_G5 | DRR432201 | Zenjimaru | G | 4 | 5 | 22,889,909 |
| 001_N1 | DRR432202 | Zenjimaru | N | 4 | 1 | 20,184,512 |
| 001_N2 | DRR432203 | Zenjimaru | N | 4 | 2 | 25,657,638 |
| 001_N3 | DRR432204 | Zenjimaru | N | 4 | 3 | 19,423,565 |
| 001_N4 | DRR432205 | Zenjimaru | N | 4 | 4 | 19,739,934 |
| 001_N5 | DRR432206 | Zenjimaru | N | 4 | 5 | 19,201,751 |
| 009_G1 | DRR432207 | Fuyuu | G | 4 | 1 | 21,613,850 |
| 009_G2 | DRR432208 | Fuyuu | G | 4 | 2 | 22,257,259 |
| 009_G3 | DRR432209 | Fuyuu | G | 4 | 3 | 20,455,356 |
| 009_G4 | DRR432210 | Fuyuu | G | 4 | 4 | 19,398,452 |
| 009_G5 | DRR432211 | Fuyuu | G | 4 | 5 | 23,050,939 |
| 009_N1 | DRR432212 | Fuyuu | N | 4 | 1 | 22,117,285 |
| 009_N2 | DRR432213 | Fuyuu | N | 4 | 2 | 25,127,318 |
| 009_N3 | DRR432214 | Fuyuu | N | 4 | 3 | 18,811,921 |
| 009_N4 | DRR432215 | Fuyuu | N | 4 | 4 | 11,076,509 |
| 009_N5 | DRR432216 | Fuyuu | N | 4 | 5 | 10,878,036 |
| 017_G1 | DRR432217 | Amayotsumizo | G | 4 | 1 | 20,607,391 |
| 017_G2 | DRR432218 | Amayotsumizo | G | 4 | 2 | 20,167,044 |
| 017_G3 | DRR432219 | Amayotsumizo | G | 4 | 3 | 21,146,098 |
| 017_G4 | DRR432220 | Amayotsumizo | G | 4 | 4 | 17,940,401 |
| 017_G5 | DRR432221 | Amayotsumizo | G | 4 | 5 | 22,367,825 |
| 017_N1 | DRR432222 | Amayotsumizo | N | 4 | 1 | 17,168,623 |
| 017_N2 | DRR432223 | Amayotsumizo | N | 4 | 2 | 7,951,916 |
| 017_N3 | DRR432224 | Amayotsumizo | N | 4 | 3 | 9,006,684 |
| 017_N4 | DRR432225 | Amayotsumizo | N | 4 | 4 | 10,138,774 |
| 017_N5 | DRR432226 | Amayotsumizo | N | 4 | 5 | 18,773,334 |
| 025_G1 | DRR432227 | Fujiwaragosho | G | 4 | 1 | 28,198,385 |
| 025_G2 | DRR432228 | Fujiwaragosho | G | 4 | 2 | 24,047,734 |
| 025_G3 | DRR432229 | Fujiwaragosho | G | 4 | 3 | 20,473,158 |
| 025_G4 | DRR432230 | Fujiwaragosho | G | 4 | 4 | 24,050,115 |
| 025_G5 | DRR432231 | Fujiwaragosho | G | 4 | 5 | 22,472,047 |
| 025_N1 | DRR432232 | Fujiwaragosho | N | 4 | 1 | 16,020,188 |
| 025_N2 | DRR432233 | Fujiwaragosho | N | 4 | 2 | 19,903,147 |
| 025_N3 | DRR432234 | Fujiwaragosho | N | 4 | 3 | 18,421,060 |
| 025_N4 | DRR432235 | Fujiwaragosho | N | 4 | 4 | 18,834,766 |
| 025_N5 | DRR432236 | Fujiwaragosho | N | 4 | 5 | 17,864,521 |
| 067_G1 | DRR432237 | Gionbou | G | 4 | 1 | 21,066,342 |
| 067_G2 | DRR432238 | Gionbou | G | 4 | 2 | 25,979,456 |
| 067_G3 | DRR432239 | Gionbou | G | 4 | 3 | 26,034,081 |
| 067_G4 | DRR432240 | Gionbou | G | 4 | 4 | 25,010,860 |
| 067_G5 | DRR432241 | Gionbou | G | 4 | 5 | 23,539,500 |
| 067_N1 | DRR432242 | Gionbou | N | 4 | 1 | 22,760,948 |
| 067_N2 | DRR432243 | Gionbou | N | 4 | 2 | 21,959,495 |
| 067_N3 | DRR432244 | Gionbou | N | 4 | 3 | 18,535,706 |
| 067_N4 | DRR432245 | Gionbou | N | 4 | 4 | 20,165,227 |
| 067_N5 | DRR432246 | Gionbou | N | 4 | 5 | 37,567,419 |
| 099_G1 | DRR432247 | Saijou | G | 4 | 1 | 20,411,952 |
| 099_G2 | DRR432248 | Saijou | G | 4 | 2 | 20,283,439 |
| 099_G3 | DRR432249 | Saijou | G | 4 | 3 | 18,866,645 |
| 099_G4 | DRR432250 | Saijou | G | 4 | 4 | 17,929,634 |
| 099_G5 | DRR432251 | Saijou | G | 4 | 5 | 19,040,690 |
| 099_N1 | DRR432252 | Saijou | N | 4 | 1 | 15,999,437 |
| 099_N2 | DRR432253 | Saijou | N | 4 | 2 | 20,414,214 |
| 099_N3 | DRR432254 | Saijou | N | 4 | 3 | 18,812,652 |
| 099_N4 | DRR432255 | Saijou | N | 4 | 4 | 21,875,433 |
| 184_G1 | DRR432256 | Iwasedo | G | 4 | 1 | 19,767,980 |
| 184_G2 | DRR432257 | Iwasedo | G | 4 | 2 | 20,100,340 |
| 184_G3 | DRR432258 | Iwasedo | G | 4 | 3 | 19,987,536 |
| 184_G4 | DRR432259 | Iwasedo | G | 4 | 4 | 21,179,999 |
| 184_G5 | DRR432260 | Iwasedo | G | 4 | 5 | 18,783,595 |
| 184_N1 | DRR432261 | Iwasedo | N | 4 | 1 | 21,808,054 |
| 184_N2 | DRR432262 | Iwasedo | N | 4 | 2 | 23,944,670 |
| 184_N3 | DRR432263 | Iwasedo | N | 4 | 3 | 15,970,404 |
| 184_N5 | DRR432264 | Iwasedo | N | 4 | 4 | 20,536,215 |
| 256_G1 | DRR432265 | Mizushimagosho | G | 4 | 1 | 21,011,950 |
| 256_G2 | DRR432266 | Mizushimagosho | G | 4 | 2 | 22,279,226 |
| 256_G3 | DRR432267 | Mizushimagosho | G | 4 | 3 | 21,709,200 |
| 256_G4 | DRR432268 | Mizushimagosho | G | 4 | 4 | 23,185,484 |
| 256_G5 | DRR432269 | Mizushimagosho | G | 4 | 5 | 21,614,892 |
| 256_N1 | DRR432270 | Mizushimagosho | N | 4 | 1 | 15,637,521 |
| 256_N2 | DRR432271 | Mizushimagosho | N | 4 | 2 | 19,052,814 |
| 256_N3 | DRR432272 | Mizushimagosho | N | 4 | 3 | 5,097,561 |
| 256_N4 | DRR432273 | Mizushimagosho | N | 4 | 4 | 29,620,807 |
| 256_N5 | DRR432274 | Mizushimagosho | N | 4 | 5 | 10,759,570 |
